# Towards modeling genome-scale knowledge in the global ocean

**DOI:** 10.1101/2023.11.23.568447

**Authors:** Antoine Régimbeau, Olivier Aumont, Chris Bowler, Lionel Guidi, George A. Jackson, Eric Karsenti, Laurent Memery, Alessandro Tagliabue, Damien Eveillard

## Abstract

Earth System Models (ESMs) highly simplify their representation of biological processes, leading to major uncertainty in climate change impacts. Despite a growing understanding of molecular networks from genomic data, describing how changing phytoplankton physiology affects the production of key metabolites remains elusive. Here we embed a genome-scale model within a state-of-the-art ESM to deliver an integrated understanding of how gradients of nutritional constraints modulate metabolic reactions and molecular physiology. Applied to the prevalent marine cyanobacteria *Prochlorococcus*, we find that glycogen and lipid storage can be understood as a consequence of acclimation to environmental gradients. Given the pressing need to assess how biological diversity influences biogeochemical functions, genome-enabled ESMs allow the quantification of the contribution of modeled organisms to the production of dissolved organic carbon and its molecular composition.

---

Earth System Models (ESMs) are a powerful tool to study the future impact of climate change on the ocean (1). However, due to computational limitations (2), they need to simplify biology and biological processes, which limits our ability to understand and constrain biological feedbacks on climate and biogeochemistry. For example, the net growth of an organism is described by a set of ordinary differential equations (3, 4) involving nutrient uptake based on a scheme introduced by Monod (5) and later extended by Droop through the use of cellular quotas (6). Following these approaches, more recent Plankton Functional Type (PFT) models (2, 7, 8) rely on extensive efforts to estimate a broad set of parameters that affect plankton functional diversity and describe traits critical to biogeochemical processes. However, this leads to a fundamental disconnect between the biological underpinnings of today’s ESMs around nutrient limitation or other phenotypical traits and the ever-growing gene- and genome-centered datasets that have emerged over recent years (8–10). Using distinct approaches, two notable modeling studies have addressed the discrepancy between molecular functions and oceanic provinces. In 2017, Coles *et al*. (9) developed a trait-based model that harnessed omics data. This approach characterized omics-derived traits, and simulated their interactions, providing a computationally feasible representation of the community’s molecular functions in an oceanic environment. More recently, Casey *et al*. (11) focused on modeling organisms, with a specific focus on the *Prochlorococcus* genus along an Atlantic transect. They achieved this by extensively parameterizing and optimizing Genome-Scale Models (GSMs) using omics data. However, this approach demands extensive data and computational power to be applied effectively at global scale. GSMs, developed primarily for bioengineering (12), offer an effective way to engage with growing biological datasets as they use gene-protein-reaction associations to represent more comprehensively the metabolic potential of an organism as defined by its genetic material (13). GSMs consider a set of reactions organized into metabolic networks in which products of some reactions become substrates for others. Its genome-scale nature requires the description of many reactions, ranging from several hundred in prokaryotes to thousands for eukaryotes. If environmental conditions are also incorporated, GSMs can predict an organism’s growth rate, the production of auxiliary metabolites, or the metabolic pathways it utilizes (14). In particular, GSMs can be employed to assess the diversity and magnitude of metabolite production that contributes to the oceanic Dissolved Organic Carbon (DOC) pool. Even though understanding individual organisms in such detail can provide crucial mechanistic insights into the structure and function of marine ecosystems (15), GSMs have never been coupled to ESMs. As a result, their potential for providing insights into global scale processes remains unknown.

To bridge these gaps, we propose a modeling compromise that balances the trade-offs around mechanistic detail and computational efficiency. Our approach aims to avoid laborious parameterization while still capturing the intricate biological complexity of organisms. In doing so, we can effectively model molecular functions in various oceanic regions without overwhelming computational demands. This compromise holds great promise for advancing our understanding of microbial processes in the vast ocean ecosystems.

## Incorporating genome-scale knowledge into biogeochemical models

Because integrating a GSM within an ESM requires the solving of several hundred equations at each grid point of the Earth, it is currently computationally unrealistic. A feasible integration needs a numerical abstraction of the dependencies between growth rate and environmental conditions. Furthermore, this abstraction must include the cellular mechanisms described by the genome (Fig. 1a.). We achieve this through a metabolic niche approach (16) that projects the entire metabolism of a species into a reduced mathematical space driven by the availability of metabolites or nutrients (Fig. 1b.,c. for illustration and see Materials and Methods for complete details). When integrated within the environmental context of ESMs, this new modeling approach benefits from recent computational advances and exploits the rapidly growing amount of omics data and associated process understanding (8–10).

**Fig. 1.**
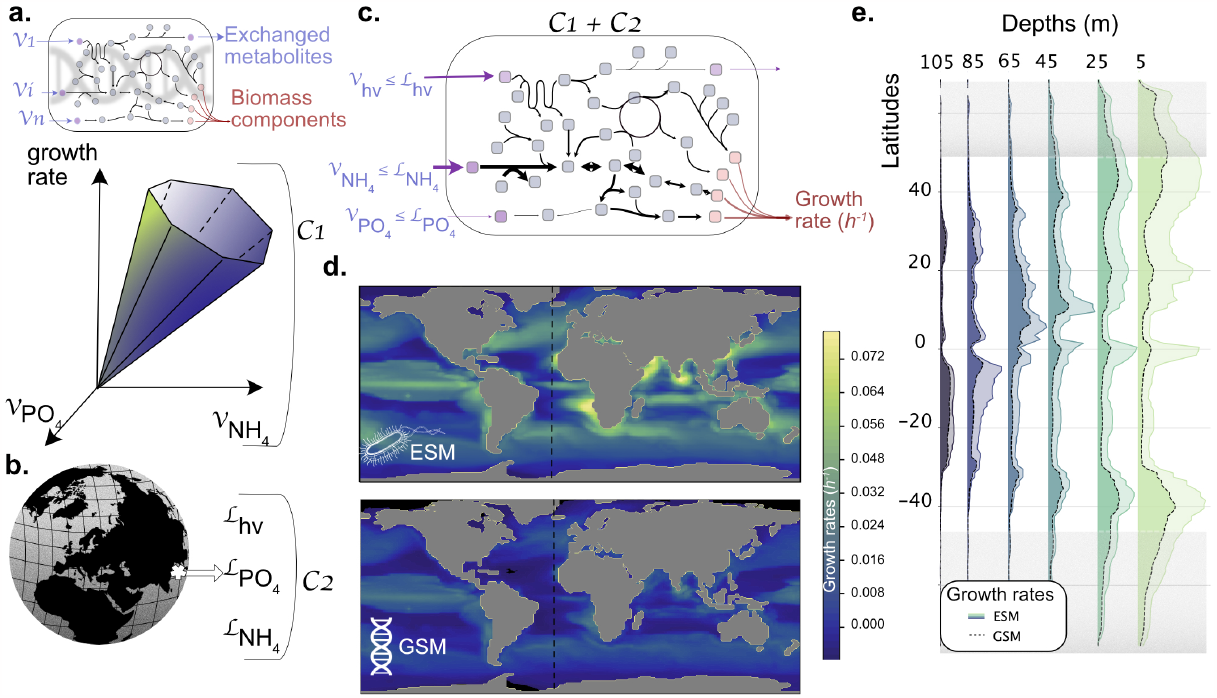
Illustration of the modeling combination between a Genome-Scale Model (GSM), i.e., *Prochlorococcus MED4* GSM, and an Earth Systems Model (ESM), i.e., NEMO-PISCES, and comparison between growth rates estimated from ESM and GSM. **a** |From a metabolic network, we defined a solution space embedding all possible fluxes that go through each network reaction. These fluxes satisfy the quasi-steady states assumption and other thermodynamic constraints defined in the GSM. This set of constraints (*C1*) is defined as biotic constraints, and they affect inner reactions, as well as exchange reactions responsible for the uptake or secretion of nutrients, and a biomass reaction simulating the growth of the organism (color scale similar to panel d., see Appendix 1 for details). **b** |Earth Systems Models predict global ocean biogeochemistry across space and time. Here the ESM provides uptake fluxes of nutrients for each grid point for each modeled organism. In our framework, these uptake values are used as a set of constraints (*C2*) on the exchange reactions of the GSM. These are defined as the set of abiotic constraints that are applied in the model. **c** |*C1* and *C2* are combined to constrain further exchanged metabolite fluxes at each grid point of the global ocean. As a result, we can estimate the organismal growth rate and all feasible inner fluxes corresponding to a given environment as proposed by the ESM. **d** |Description of growth rates (*h*^*−*1^) at 5 m depth estimated from NEMO-PISCES picophytoplankton (top) and *Prochlorococcus MED4* GSM (bottom). The dashed line shows the transect described in the following panel. **e** |Distribution of respective growth rates across latitudes and depths at longitude -24°. Grey areas indicate latitudes that do not allow *Prochlorococcus MED4* growth because of thermal limits; the GSM does not consider them. The relationship between growth rates across space (above 500 m) and time (i.e., without gray areas) shows R^2^:0.80 and slope: 0.787 (see Extended Fig. 11).

In this work, we combine a GSM with the quota version of the NEMO-PISCES global ocean biogeochemical model (17), which is a classic example of a coupled ocean physicochemical-biological model embedded within an ESM used for climate change studies. NEMOPISCES predicts the spatiotemporal distribution of three coarse-grained cosmopolitan phytoplankton groups (picophytoplankton, nanophytoplankton, and diatoms) considering various environmental conditions such as temperature, light, and a range of major and macro nutrients (Fig. 1b). NEMO-PISCES estimates the environmental conditions and resulting growth rate for each of the three phytoplankton groups. In this novel approach, we employ the same set of conditions to calculate offline the growth rate using a selected GSM over the annual cycle (Fig. 1c; see Materials and Methods for more details). To appraise our approach, we compared the growth rate of the picophytoplankton group simulated by NEMO-PISCES with the growth rate calculated using the specific GSM of *Prochlorococcus MED4* (18) (Fig.1d,e). Encouragingly, although *Prochlorococcus MED4* represents only one specific strain of picophytoplankton, simulations using its GSM were able to qualitatively reproduce the patterns of average monthly NEMO-PISCES picophytoplankton growth rates both at the surface (Fig. 1d) and at different depths (Fig. 1e) over the global ocean (*r >* 0.9; Extended Fig. 11). Similar prediction abilities were found for two groups of diatoms (Extended Fig. 12), implying the potential ability to assess the result of competition between the two diatom species that have GSMs available *Thalassiosira pseudonana* (19) and *Phaeodactylum tricornutum* (20) in conditions computed by NEMO-PISCES for diatoms (Extended Fig. 13). While qualitatively similar to NEMO-PISCES, the predictions show an expected quantitative mismatch. A key difference is that NEMO-PISCES models the entire community of different phytoplankton functional groups (picophytoplankton, nanophytoplankton, and diatoms), whereas a GSM represents only one strain within these communities. To better quantitatively align the two models, it is necessary to incorporate GSMs that embed diverse ecotypes (21, 22), as observed with *in situ* data. For instance, the abundance of the MED4 strain represents only about one-third of the total *Prochlorococcus* abundance (21). Furthermore, despite being driven by flux estimates, GSM predictions exhibit qualitative alignment with *in situ* abundance and concentration patterns (see Appendix 4.D), allowing us to explore the physiological implications further.

### GSM-based predictions of *Prochlorococcus MED4* in the global ocean

Unlike ESMs, which require new parameterizations and a set of parameter values for each newly introduced trait, GSMs can reveal any flux occurring through a metabolic process within an organism at the intracellular level as long as it is defined within the metabolic network. Our simulations are thus not limited to growth rate estimates, but can be exploited to quantify production of any metabolite represented in the GSM, including primary and secondary compounds, and to investigate the activity of the corresponding pathways in response to environmental gradients. Based on more than 10^6^ environmental conditions provided by NEMO-PISCES across space and time over a year, we estimated for each condition the growth and potential metabolic content of *Prochlorococcus MED4*.

Among the different types of carbon storage available in *Prochlorococcus MED4* GSM, we observed that pyruvate and glycogen productions were fully correlated (*r* = 1). In contrast, glycogen and lipid productions show a lower correlation (*r* = 0.7). While the production of lipids and glycogen by *Prochlorococcus MED4* shows similar qualitative patterns across the surface ocean, except in tropical Atlantic regions and the Bay of Bengal (Extended Fig. 10), they differ across depths and latitudes (Fig. 3a). This insight is only possible because the information about such traits is accessible from GSMs. Our model explains these two strategies by the parallel variation in a so-called ‘resource constraint’. Unlike previous work that focuses on statistical description (24) or proteomic measurements (25) of resource limitation, the resource constraint we use here results from the GSM and represents how the nutrient uptake variability affects the organism’s growth. For a given nutrient, a low resource constraint indicates that a high quantity of this nutrient can be used for processes not related to growth, like secondary metabolite production. Thus, a low resource constraint implies relative growth stability under variations in nutrient bioavailability. In contrast, a high resource constraint implies that only a small amount of the nutrient can be allocated outside growth, making growth more sensitive to its variation and reducing its availability for other processes, such as secondary metabolite production. At its maximum (100%), the resource constraint indicates that the given nutrient limits the organism’s growth. Using this definition, our model predicts that the ocean surface is not limited by light as expected (i.e., low light constraint). Moreover, nitrogen and phosphorus emerge at higher resource constraints in distinct zones (Fig. 2a). Specifically, growth of*Prochlorococcus MED4* is limited by phosphorus in the central Atlantic and the Indian Oceans (Fig. 2c). This aligns with the quota estimation of NEMO-PISCES, which predicts a lower P:N ratio in these areas (Fig. 2b). Nitrogen-limited provinces are in the Southern and North Atlantic Ocean, which show an antagonistic pattern between nitrogen and phosphorus resource constraints. We summarised the above results in a Principal Component Analysis (Fig. 2a), which highlights a positive correlation between growth rate, glycogen, lipid content, and Photosynthetically Active Radiation (PAR), suggesting acclimation strategies that require further investigations.

**Fig. 2.**
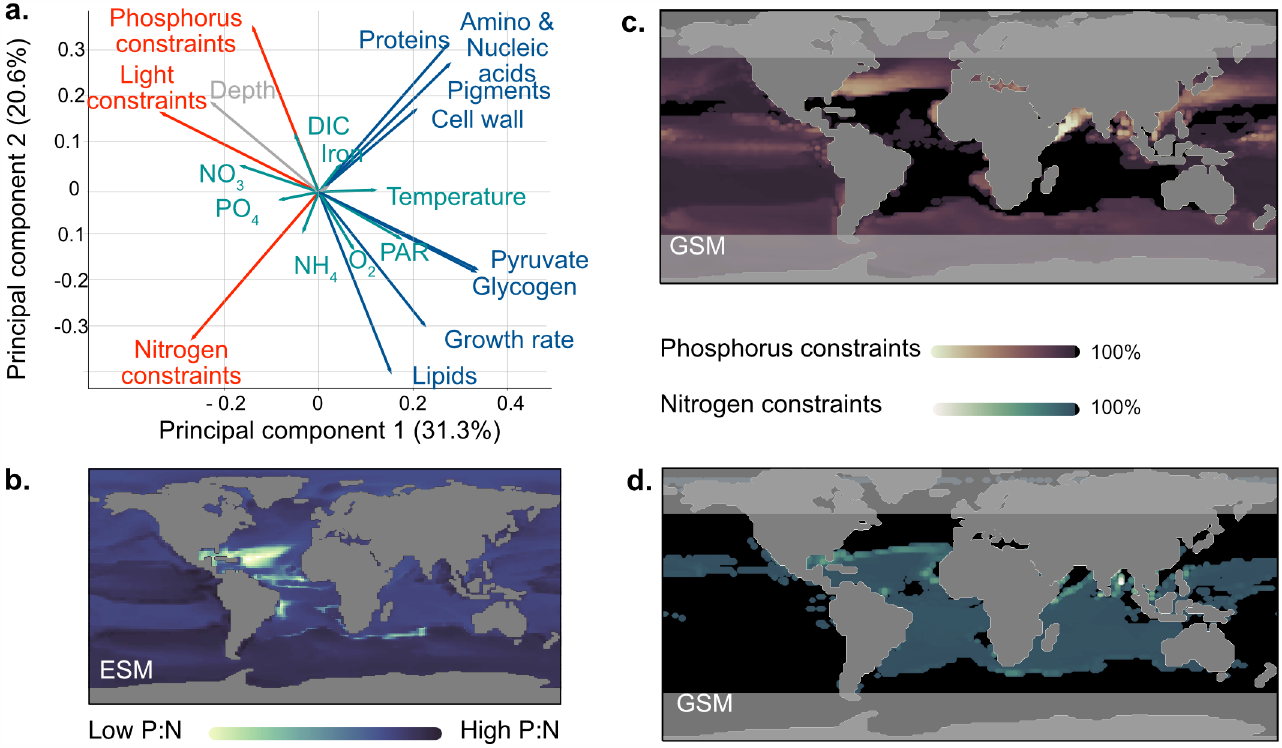
Simulation of *Prochlorococcus MED4* genome-scale model across the global ocean. **a** |Principal component analysis of over 10^6^ predictions in the global ocean, across depths and over a year. Physiological factors emphasized by GSM modeling are blue for organismal composition and red for resource constraints, whereas environmental factors are in green and those associated with biogeography in grey. **b** |Distribution of the phosphorus to nitrogen ratio estimated by NEMO-PISCES for picophytoplankton. **c** |Distribution of phosphorus constraints based on orthophosphate uptake fluxes at the surface ocean in January. Lighter colors indicate no resource constraints on uptake. In contrast, higher resource constraints are depicted with darker colors until their maximum (100% described in black) when the nutrient is limiting per se. **d** |Distribution of nitrogen constraints based on ammonium uptake fluxes (the only source of nitrogen available to *Prochlorococcus MED4* (23)) following the same color nomenclature as in panel **c** .

**Fig. 3.**
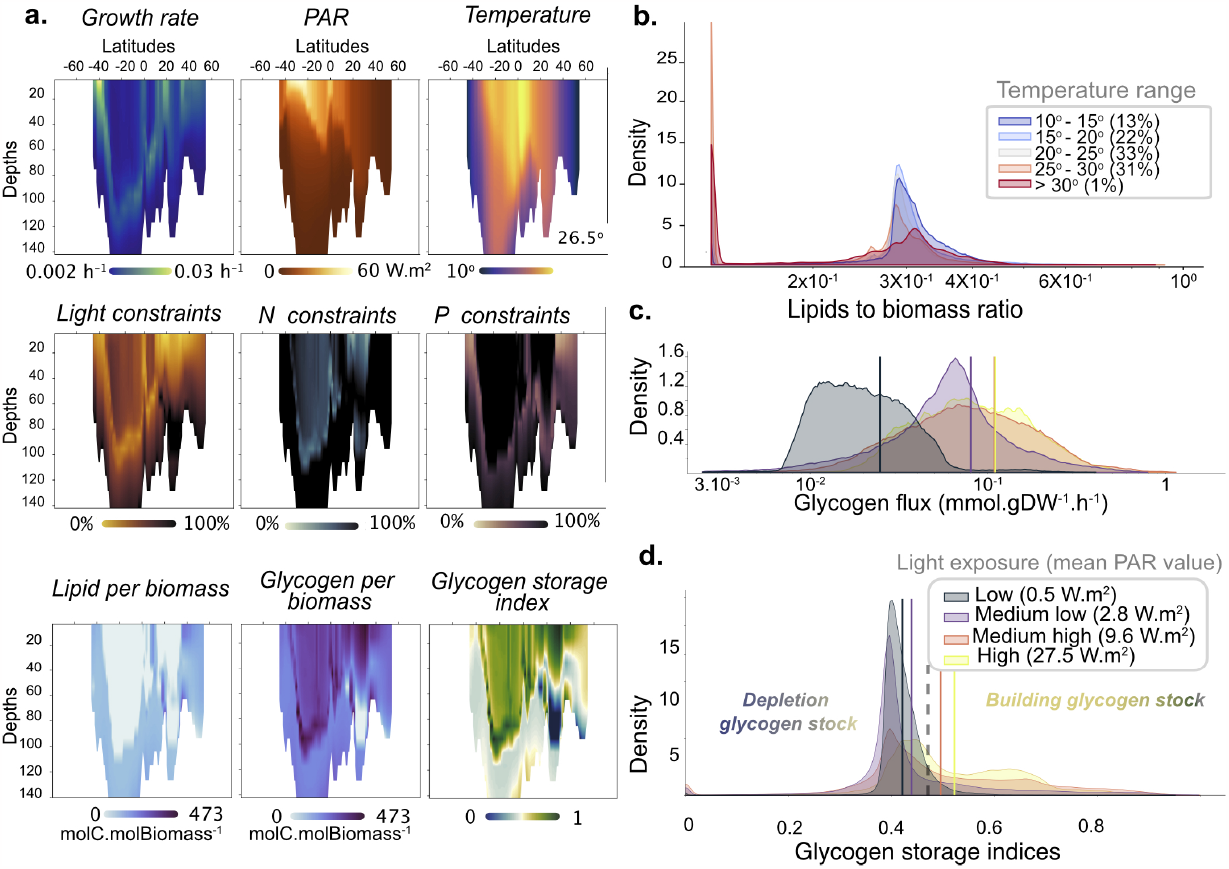
Investigation of *Prochlorococcus MED4* genome-scale model fitness and acclimation strategies across the global ocean (more than 10^6^ estimations). **a** |Description of *Prochlorococcus MED4* genome-scale behavior across the Atlantic Ocean transect (longitude -24° ). It describes growth rate according to light and temperature, associated nutrient constraints (light, nitrogen, and phosphorus), and acclimation consequences (glycogen and lipid productions per biomass) resumed by the glycogen storage index (see Material and Methods for details), with blue colors indicating consumption of potential glycogen stock and green colors showing increased storage. **b** |Distribution of lipid contribution to *Prochlorococcus MED4* biomass over five distinct temperature ranges (see Materials and Methods for details). **c** |Distribution of glycogen production satisfying predicted *Prochlorococcus MED4* growth rates under four categories of gradual light exposures. **d** |Distribution of glycogen storage indices computed with estimated *Prochlorococcus MED4* growth rates under four similar categories of gradual light exposures. Indices between 0 and 0.46 index indicate a gradual decrease of glycogen stocks to support growth. Above 0.46, indices are associated with full phototrophic growth with increased glycogen storage.

### GSM-based acclimation strategies of *Prochlorococcus MED4* in the global ocean

Linear relationships between specific growth rate and growth requirements are too restrictive to capture the underlying acclimation strategies deployed by organisms. For instance, along ocean transects, simulated data representing *Prochlorococcus MED4* growth rates, light, and temperature are not linearly related (Fig. 3a). Patterns in resource constraints better explain *Prochlorococcus MED4* growth. Acclimation to these constraints and environmental parameters, such as temperature, produce distinct carbon storage strategies. Either carbon is stored in the form of lipids or as glycogen. Their respective production rates can thus reveal the prevalent acclimation strategy of *MED4*.

Overall, provinces with high nitrogen constrained growth exhibit similar lipid and glycogen production patterns. When limited by phosphorus, *Prochlorococcus MED4* shows more carbon storage in the form of glycogen than lipids (Fig. 3a). This can be explained by the fact that the lipids used here are a mix of different types, including phospholipids. Lipids are generally produced in extreme conditions, such as at depth and at high latitudes, far from the organism’s optimal growth conditions. By studying the distribution of *Prochlorococcus MED4* lipid to biomass ratios among all possible environmental conditions, we found that it can increase three-fold in cold waters (relative to *<* 25*°C* conditions; Fig. 3b), consistent with molecular evidence from other cyanobacteria (26). When less constrained by light, *Prochlorococcus MED4* growth is associated with the production of carbon compounds that are metabolically faster to access, such as glycogen. High glycogen production is observed when carbon is minimally stored as lipids. Moreover, when both types of stocks can be used, the observed difference in their production rates is due to the higher energy needed to produce lipids compared to glycogen. When grouped into four categories of increasing light exposure, the mean value of glycogen production per category increases (Fig. 3c), linking this process to a photosynthetic behavior. In this respect, by investigating the inner machinery of *Prochlorococcus MED4*, we can quantify the amount of carbon used for biomass or glycogen production. We defined the glycogen storage index as the normalized ratio of carbon allocated to glycogen production over the total amount of carbon fixed. This index represents the ability of the organism to store glycogen. A high index (i.e., 1) indicates high use of carbon for glycogen storage while growing at maximal capacity. In contrast, a lower index (i.e., 0) reveals a lack of glycogen production and the channeling of carbon toward growth.

In the GSM, lack of production and consumption are interrelated. Below the mean glycogen storage index value (i.e., 0.46), *Prochlorococcus MED4* combines photosynthesis and glycogen consumption (or lack of production) to ensure a higher growth rate, whereas, above this value, it shows a growth regime with glycogen storage or secretion. Our indices show a natural tendency for the organism to undergo glycogen consumption in low-light conditions (Fig. 3d). Conversely, MED4 displays growth and glycogen over-production in regions where phosphorus constrains growth rate, which emphasizes the importance of estimating the nitrogen and phosphorus constraints to assess growth regime and to uncover long-term vs. short-term carbon storage strategies (i.e., lipids vs. glycogen). These results highlight the importance of examining the behavior of GSMs in sub-optimal conditions to assess different acclimation strategies more deeply. Furthermore, it reveals how the high level of metabolic flexibility impacts biomass composition in the global ocean and indicates which traits should be incorporated into ESMs.

### Predicting hot spots of biotic production and metabolite diversity

By assessing genome-scale knowledge, GSMs are emerging as valuable tools for investigating cellular composition (for detailed information and its application in designing new trait models, refer to Supplementary Material 5.A) and diverse metabolite contents. As a case in point, specific metabolites play crucial roles in the labile Dissolved Organic Carbon (DOC) pool, a fundamental component of the ocean carbon cycle (27), and are significant in bacterial and plankton growth (28). Surprisingly, with few exceptions(29), current ESMs overlook this diverse range of metabolites when modeling DOC and instead represent a generic DOC pool. To address this shortcoming, we compiled a comprehensive summary of the DOC metabolite compounds listed in (27), specifically focusing on those produced by *Thalassiosira pseudonana* and *Prochloroccocus MED4* GSMs. Our analysis revealed that *T. pseudonana* and *Prochloroccocus MED4* GSMs produced 33 and 19 metabolites, respectively (see Appendix Table.1.D for detailed information). To estimate the contribution of each GSM to DOC production, we aggregated all the metabolite flux estimates (Fig.4; see Materials & Methods for details). We furthermore investigated the diversity and abundances of metabolites supporting DOC flux production in terms of seasonal variation across each GSM. As anticipated, upwellings and fronts demonstrated higher DOC production and a wider array of secreted metabolites. Interestingly, the analysis using GSMs shows opposite patterns in DOC production and metabolite diversity. Despite producing more diverse metabolites, *Prochloroccocus MED4* shows restricted areas of high metabolite diversity amidst wider arrays of high DOC production. On the contrary, *T. pseudonana* displays higher DOC production amidst wider areas of high diversity. By comparing both GSMs, *T. pseudonana* played a predominant role in the magnitude of DOC production and the diversity of associated metabolites. The more expansive areas of DOC diversity are consistent with the importance of diatoms in determining the biogeography of bacterial heterotrophs (30). Through a broader analysis of provinces, we distinguished regions driven predominantly by diatom influences from those more strongly affected by *Prochloroccocus*, reaffirming earlier findings (31). Notably, provinces exhibiting high metabolite diversity did not necessarily align with high DOC production, highlighting the importance of further investigation to understand the support for trophic interactions via metabolic cross-feeding between organisms (32) or to improve predictions concerning the relationship between functional diversity and ecosystem productivity (28, 33).

**Fig. 4.**
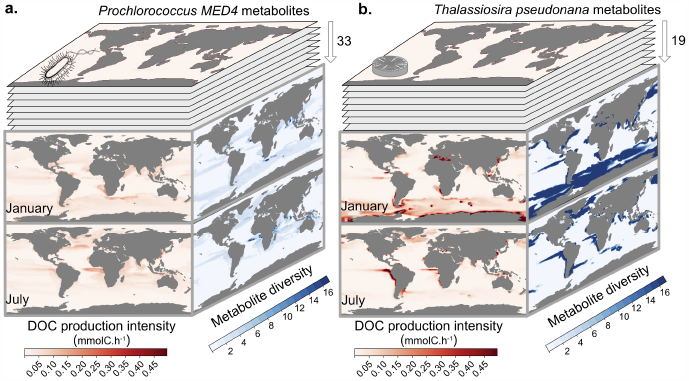
Contribution of Genome-Scale Model to Dissolved Organic Carbon (DOC). **a** |*Prochloroccocus MED4* produces 33 metabolites that contribute to DOC pool. The compilation of all compounds allows for an estimation of the intensity of DOC production in red, in January (top layer) and July (bottom layer). It also allows estimation of the diversity of metabolites involved in the DOC production in blue in January and July. **b** |*Thalassiosira pseudonana* produces 19 metabolites associated with the DOC pool. A similar analysis as in panel a. identifies hot spots for DOC production intensity and associated metabolic diversity in January and July.

## Discussion

Our findings underscore the remarkable potential of integrating genomic knowledge into ESMs. This conceptual convergence was expected (27), but achieving it has been hindered by the challenges of extensive data integration and costly measurements (9, 11). By incorporating GSMs into a biogeochemical model, we can estimate growth rates and assess cellular composition, which partially aligns with observations. While our modeling paradigm represents a significant step forward, there is room for refinement, particularly in improving GSMs to accurately capture *in situ* and diurnal growth rates. Nevertheless, this approach allows for predicting complex and diverse molecules, such as cryptic metabolites (e.g., DMSP production in Extended Fig. 13). The combined understanding of these metabolites contributes to our comprehension of dissolved organic carbon (DOC) production. GSMs enable estimates of DOC production for each modeled organism and describe each molecular compound associated with it together with their relative proportions. However, these new estimates, driven by omics-derived information, must be further validated against the next generation of quantitative molecular data (34, 35), necessitating mesoscale studies to refine our modeling efforts.

In perspective, our modeling framework is ready for incorporating recent genome-scale resources, such as the reconstruction of metagenome-assembled genomes (MAGs) (36), for a more accurate representation of biodiversity (37) and its implications for complex molecule production. Furthermore, incorporating metabolic frameworks into ESMs opens opportunities to explore advanced evolutionary theories involving gene transfer or modulation at global scales (38).

Given the pressing need to understand how biological diversity influences global biogeochemical functions, our mathematical framework serves as an important bridge to better connect the concluion of intergovernmental panels addressing climate change and biodiversity loss (IPCC and IPBES respectively). The integration of these cutting-edge approaches promises to advance our understanding of Earth’s intricate microbial ecosystems and their impact on global biogeochemical processes in a changing ocean.

## Material and methods

### Genome-Scale Model

A Genome-Scale Model (GSM) is stated as a set of linear constraints, representing the quasi-steady state assumption, and the thermodynamic constraints:

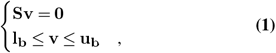

The matrix **S** ∈ ℝ^*n,m*^ abstracts the metabolic network of *n* metabolites and *m* reactions, the vector **v** ∈ ℝ^*m*^ represents the fluxes that go through each network reaction, **l**_**b**_, **u**_**b**_ ∈ ℝ^*m*^ are the lower and upper bounds of **v**. To represent the organism’s growth rate, metabolic models include a biomass reaction that describes the metabolic requirement for an organism to grow. It is included in **S** and cannot have a negative flux. Given the stated problem, one can calculate with a dedicated solver and extract the flux for each network reaction, including the biomass reaction. The solution is one of the feasible physiological states of the system. In this state, one can estimate the organism’s growth rate as the flux through the biomass reaction. For more detail on the metabolic framework and GSM, the reader is referred to Appendix 1.

### Metabolic niche projection

We call *solution space* ℱ the convex hypervolume composed of **v** satisfying Eq. 1. ℱ is defined in a space where each dimension represents the flux through a reaction. Investigation of this space is subject to numerous techniques in the context of metabolic engineering (see Price et al. 2004 (39) for review). However, using ℱ is not well suited in biogeochemical models because of its size and complexity. In most cases, one cannot describe the entire shape of ℱ as its complexity grows exponentially with the number of reactions in the GSM. In addition, biogeochemical models describe the distribution of a few nutrients compared to the number of metabolites in a GSM. It means that most of the reactions and underlying mechanisms of the metabolic network can be abstracted in favor of a numerical tool linking the exchange reactions relative to nutrients available in biogeochemical models with the biomass reaction to estimate a growth rate. In mathematical terms, it implies projecting ℱ onto a smaller space composed of the sole reactions of interest (i.e., reactions representing a parameter in the biogeochemical model and the biomass reaction). This projection, called the metabolic niche, is computed through Multi-Objective Linear Programming (40). Decomposing the flux vector 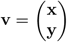 where **x** is a flux vector composed of the reactions of interest (i.e., the exchange reactions concerning the nutrient and light absorption) and **y** is a flux vector composed of all the other reactions. The projection of ℱ is equivalent to solving the following problem :

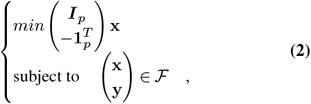

where **I**_*p*_ is the identity matrix in ℝ^*p×p*^, **1**_*p*_ is the column vector composed of ones in ℝ^*p*^, and 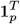 its transposition. The solution of 2 without its last component gives a set of vertices describing the new convex hypervolume 𝒩. Applied to several organisms, the investigation of the metabolic niche hypervolume has shown ecological properties (16).

### Description of the biogeochemical model

Our framework is interoperable with the marine biogeochemical model PISCES (Pelagic Interactions Scheme for Carbon and Ecosystem Studies). PISCES is the biogeochemical component of the NEMO (Nucleus for European Modelling of the Ocean) modeling platform. This study uses the quota version of PISCES (17). Three phytoplankton groups are explicitly modeled (picophytoplankton, nanophytoplankton and diatoms) whose growth rates are limited by iron, phosphate, nitrate, ammonium and silicate availability. Two zooplankton groups (micro- and mesozooplankton) are represented. PISCES also models dissolved oxygen, particulate and dissolved organic matter, and calcite. The uptake of nutrients and phytoplankton growth rates are modeled using quota formalism. Metabolic rates increase with temperature according to the commonly used Eppley parameterization (41). Based on the environmental conditions and the biotic interactions between the different plankton groups, NEMO-PISCES estimates the growth rate for each plankton group using partial differential equations.

### The metabolic niche in NEMO-PISCES

Thanks to the metabolic niche projection, we can compute the growth rate of an organism based on the environmental conditions dictated by NEMO-PISCES. Thus, biology will be handled by omic-derived knowledge of metabolic models, while NEMO-PISCES will compute the nutrient availability. NEMO-PISCES’s simulations provide nutrient inputs for the algae consisting of iron, nitrate, ammonium, phosphate, silicate (used only by diatoms), and the quantity of carbon fixed by photosynthesis. These fluxes are also used in distinct metabolic niches of GSMs representative of these generic phytoplanktons. *Prochlorococcus* MED4 cannot assimilate nitrate but does assimilate ammonium, iron, and phosphate. NEMO-PISCES inputs constrain the exchange reactions of the previous metabolite and the 3-phospho-D-glycerate carboxylase reaction for the quantity of carbon fixed. However, our calculations did not incorporate iron (see Appendix 5.B). In the context of generic diatom modeling (*Thalassiosira pseudonana* and *Phaeodactylum tricornutum*), their GSMs (18, 19) did not consider iron. The equivalent reaction for the carbon-fixed quantity is the carboxylation of ribulose-1,5bisphosphate, called *RUBISC_h* in both models. Adding to those reactions, the biomass reaction estimates a growth rate, and we have the reactions of interest that will compose the metabolic niche of each organism.

Worth noting, *Prochlorococcus* MED4’s GSM predicts growth rates outside of its thermal range (Fig.1e gray areas), as the modeling paradigm does not incorporate the thermal tolerance of *Prochloroccocus* MED4 (i.e., 10°*C*), indicating that the sole metabolism does not include processes involved in the thermal tolerance. This absence of thermal effect in the GSM does not change the overall results, as the temperature is modeled in NEMO-PISCES (41), and impacts the uptake fluxes of nutrients.

### Growth rate extraction from the metabolic niche

The metabolic niche describes the ability of an organism to survive and grow considering its genome-scale metabolic description under particular environmental conditions, here a set of nutrient concentrations, including light which is treated as a nutrient through the quantity carbon fixed by the organism. We will use this metabolic niche to obtain the maximal growth allowed by the model in the environmental conditions computed by NEMO-PISCES. Formally, considering a vector of uptake fluxes given by the biogeochemical model **x**_*envb*_, we construct 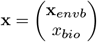, where *x*_*bio*_∈ ℝ^+^ is the flux through the biomass reaction. Then, we need to look for the maximal *x*_*bio*_ that satisfy **x** ∈ 𝒩 for a particular **x**_*envb*_ :

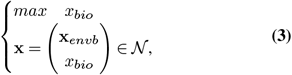

where 𝒩 is the metabolic niche described above. There are two cases of this problem. Either a solution exists, and we can solve the problem and output the solution, or there is no solution, meaning that the environmental condition does not belong to the niche. In this case, the organism cannot grow, and the growth rate should be fixed to 0. However, instead of regarding **x**_*envb*_ as a fixed nutrient uptake, we can view it as the bioavailability of nutrients. In this context, nutrient bioavailability does not represent the actual uptake of the organism; rather, it represents the upper limit of nutrient uptake. In other words, the organism is unable to take up more nutrients than what is available.

### Nutrient bioavailability from the NEMO-PISCES biogeochemical model

Indeed, the organism is not necessarily using all the resources of its environment. The metabolic network should handle the quantity of nutrients it consumes. If we denote **x**_*envb*_ as the quantity of bioavailable nutrients, and **x**_*env*_ as the actual nutrient fluxes used by the model, we need to assure that **x**_*env*_≤ **x**_*envb*_. We depict this as an additional constraint on the uptake fluxes, which changes the formulation of Eq.3. Thus, we seek for the maximum of *x*_*bio*_ that satisfies **x**_*env*_ ≤**x**_*envb*_ and 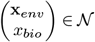. The new formulation is, therefore:

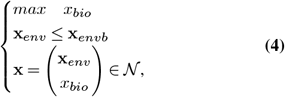

where **x**_*env*_ is the actual uptake fluxes used by the GSM and constraint by **x**_*envb*_ the uptake fluxes computed by NEMOPISCES. This new formulation assures a solution to the problem.

### Auxiliary flux computation

Not only can the metabolic niche produce growth rate estimates, but it can also estimate fluxes through any reaction of the GSM. Indeed, one can compute the metabolic niche with one additional dimension and analyze the flux variability on this dimension. Taking the former formalism for **x** we can write 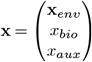, where *x*_*aux*_ ∈ ℝ is the flux through another reaction of the network, say the flux through the exchange reaction of DMSP, or the modeling reaction producing the organism pigment. With the previous method, we can determine the maximal *x*_*bio*_ with respect to **x**_*env*_, which give us 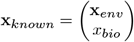. Applying the same computation on 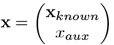 gives us a range of flux under the environmental condition defined by PISCES and the assumption that the organism is maximizing its growth rate. Rewriting Eq. 4, with *x*_*aux*_ and computing not only the maximum value but also its minimum, we have the following problem:

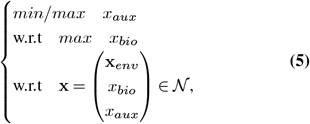

Once solved, it gives us the flux range of the new reaction of interest. This method can be applied to internal or exchange reactions.

### Component of the cellular composition

While the biomass reaction of *Prochloroccocus* MED4’s GSM has a fixed stoichiometry, we can add an exchange reaction for the components of interest to allow secretion of those components. In other words, adding a new exchange reaction allows the organism to over-produce the component. This overproduction can be estimated through the auxiliary flux computation, with an additional constraint: the model cannot uptake the component, only produce it (See Appendix 1.B for details).

### Carbon cycle hot spots

Carbon hot spots were identified using the auxiliary fluxes computation as described above. Metabolites used for the computation are the same as those described in (27) and found in the GSM of *Thalassiosira* or *Prochloroccocus* with an exchange reaction. Each flux was scaled with the carbon content of the corresponding metabolites (see Appendix **??** for detail), its unit is thus mmolC.gDW^*−*1^.h^*−*1^. To generate the intensity of DOC production at each grid point, we multiply the highest flux value by the abundance of the organism, giving us a flux in mmolC.h^*−*1^. Worth noticing, the abundance used was the one computed by NEMO-PISCES, that is, the entire diatom abundance for *Thalassiosira*, and the entire picophytoplankton abundance for *Prochloroccocus*. We included a metabolite in the diversity score if its production was above 5% of the maximum flux computed among all other metabolites.

### Resource constraint estimate

Our results show that when an organism is under the limitation of one nutrient, the others are in excess. In short, the resource constraint represents the quantity of nutrient that can be allocated to other metabolic pathways than those linked to biomass production. Our resource constraint definition is proportional to the amount of nutrient that is in excess. We can write the resource constraint on the nutrient *n* as *RC*_*n*_ ∼ −*δn* where *δn* is the quantity of the nutrient *n* that the organism can use for something other than its growth. Hence a high resource constraint means that the nutrient almost limits the production of biomass (*δn*∼ 0 ). In contrast, a low resource constraint means more nutrient *n* can be used for other products such as energy storage or other organic compounds secretion.

Formally we first consider the distance **d** between 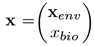 and 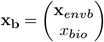 where *x*_*bio*_ is solution of Eq. 4. **x**_*env*_ is the quantity of nutrients used by the model to produce *x*_*bio*_ of growth. Whereas **x**_*envb*_ is the bioavailability of nutrients. Each component of the computed distance is a quantity of nutrient not used by the model. As this distance is defined for each environement (*env* ∈ ε where ε is the ensemble of environmental conditions and *env* is one environmental condition), we then normalize the distribution of each component *n* corresponding to one nutrient, to get a value between 0 and 100%.

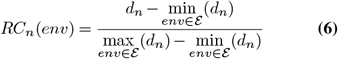

### Glycogen Storage Index

Our Glycogen Storage Index is based on the production of glycogen and the quantity of carbon fixed by the organism. Formally we write 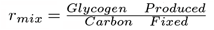 as the Glycogen Storage ratio that we normalize to give our index. ‘*Glycogen Produced*’ is the flux of glycogen for a given condition and a growth rate. ‘*Carbon F ixed*’ is the quantity of carbon fixed, provided by NEMO-PISCES. This ratio represents the quantity of carbon fixed used to produce glycogen. In other words, it looks at the resource allocation of the organism. The storage or secretion of glycogen can be used by *Prochlorococcus MED4* to adapt to different environmental conditions. From the mean value of the index, we distinguish two types of growth. Suppose the ratio is higher than its mean value. In that case, the organism is already growing at its full potential considering its environment and can store the excess carbon into glycogen. On the other hand, an index below the mean indicates that more carbon fixed is used for the biomass, and the lack of glycogen produced can be seen as consumption: the difference between the mean index and the current index is proportional to the quantity of glycogen needed by the organism to grow in a particular environment.

### Limitations of the model

Our framework, like any modeling approach, has certain limitations (see Appendix 4.D for more details). In our model, light is represented by the quantity of carbon fixed, which organisms can utilize for growth or other metabolic processes. However, organisms do not have the ability to choose not to utilize light; instead, they adapt their composition to absorb varying amounts of photons through photoadaptation and photoinhibition mechanisms. Unfortunately, these mechanisms are not accounted for in the GSM, which explains the results pertaining to pigment production (see Appendix 5). Furthermore, the quantity of carbon fixed relies on a parameter *α* (photosynthetic efficiency), which assumes uniformity across all organisms, despite experimental evidence indicating variations in these parameters (42).

The biomass reaction in our model approximates the growth rate of the GSM. However, this reaction is constructed based on laboratory experiments that may not fully capture *in situ* environmental conditions. For example, while iron is present in the *Prochlorococcus MED4* GSM, its utilization is not possible due to stoichiometric differences compared to NEMOPISCES (see Appendix 5.B).

Lastly, it is important to note that all simulations are conducted offline. Initially, NEMO-PISCES is executed, and subsequently, growth is diagnosed using the GSMs. Further work is necessary to fully integrate the GSMs into NEMOPISCES.

## Data availability

All data and codes are available on a private cloud (link in Appendix 3). The data and codes will be available behind a specific DOI upon acceptance.

## Acknowledgments

We thank the commitment of the following sponsors: CNRS (in particular the Research Federation for the study of Global Ocean Systems Ecology and Evolution, FR2022/Tara OceansGOSEE), the H2020 project AtlantECO (award number 862923), the CNRS 80 Prime grant Hoummus, the ANR program APERO (ANR-21-CE01-0027), and CINNAMON (ANR-17-CE02-0014). Computational support was provided by the bioinformatics core facility of Nantes (BiRD Biogenouest), Nantes Université, France. This article is contribution number XXX of Tara Oceans.

## Supplementary Note 1: Genome-scale model formalism and its manipulation

### 1.A. Genome scale model

From the genome or the proteome of an organism, one can associate reactions representing the metabolic abilities of an organism^1^. Some of those reactions make reactants for others, with this set of reactions denoted as a metabolic network. From this network, one can build a Genome-Scale Model (GSM) that focuses on the flux carried by each reaction under some constraints. We differentiate two types of reaction in a GSM: the internal reactions involving metabolites that represent the organism machinery and the exchange reactions involving external and internal metabolites that describe the exchanges between the organism and its environment. For a reaction *R*_*i*_, we define the stoichiometric coefficient for each internal metabolite *j* by:

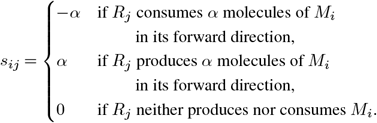

By convention, exchange reactions are written as:

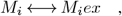

where *M*_*i*_*ex* is the external metabolite *M*_*i*_. Hence, the forward reaction means that the system adds *M*_*i*_ into the environment, whereas a reverse reaction means that the system consumes *M*_*i*_ from the environment. Consider a metabolic network of *n* reactions and *m* internal metabolites. According to kinetic theory, the change over time of the concentration of the metabolite *i* is given by the mass balance equation:

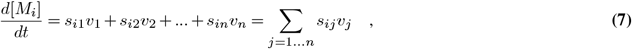

where *v*_*j*_∈ ℝ is the reaction rate or flux associated with reaction *R*_*j*_ and *s*_*ij*_ are the stoichiometric coefficient described above. Fluxes here are expressed as a mole of product formed (or mole of reactant consumed) per gram of dry weight of the considered organism per hour, i.e., mol.gDW^*−*1^.h^*−*1^. We can write the above equation for all internal metabolites expressed in vector notation as:

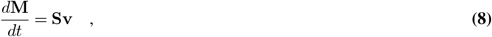

where **S** ∈ ℝ^(*n,m*)^ is called the stoichiometric matrix of the network, **v** ∈ ℝ^*n*^ the flux vector representing the flux carried by each reaction, and **M** ∈ ℝ^+*n*^ the vector composed of each metabolite concentration [*M*_*i*_]. Based on the principle that environmental changes are very slow compared to metabolic adjustments ^2^, one can assume the system at quasi-steady-states, linearising the above equations to:

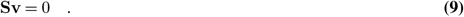

Kinetic parametrization of reactions is not well suited to our framework as it is computationally very demanding and needs extensive kinetic data to estimate enzymatic activities ^3^. Instead, GSM uses bounds, representing the thermodynamic feasibility of the reaction. A reaction cannot have an infinite flux. Thus each *v*_*i*_ are constraint as follow:

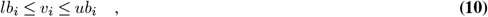

where *ub*_*i*_ represents the upper bound of the flux, meaning the highest rate of the direct reaction, and *lb*_*i*_ represents the lower bound of the flux, i.e., the highest rate of the reverse reaction. Moreover, the irreversibility of the reaction can be translated into thermodynamic constraint. For instance, a reaction known to be direct and irreversible will have a positive flux:

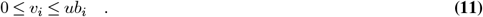

Hence the GSM is stated as a set of linear constraints:

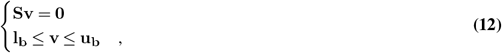

To represent the growth rate of the organism, metabolic models include a biomass reaction that describes the metabolic requirement for an organism to grow. It is included in the matrix **S** and cannot have a negative flux. Given the stated problem, one can calculate with a dedicated solver and extract the flux for each network reaction, including the biomass reaction. The solution is one of the feasible physiological states of the system. In this state, one can estimate the organism’s growth rate as the flux through the biomass reaction.

### 1.B. Growth rate and biomass components

Most models are provided with a biomass objective function in the form of a synthetic reaction. This reaction encloses the metabolic need for the organism to grow. In the case of *Prochloroccocus MED4*, the biomass reaction requires several metabolites (see Fig 5) ^4^. All those compounds have a fixed stoichiometry in the model and allow biomass production, i.e., the organism’s growth. Most of the fictive metabolites have the only purpose of meeting the metabolite requirement for biomass production. Hence, they are not consumed elsewhere.

**Extended Fig 5.**
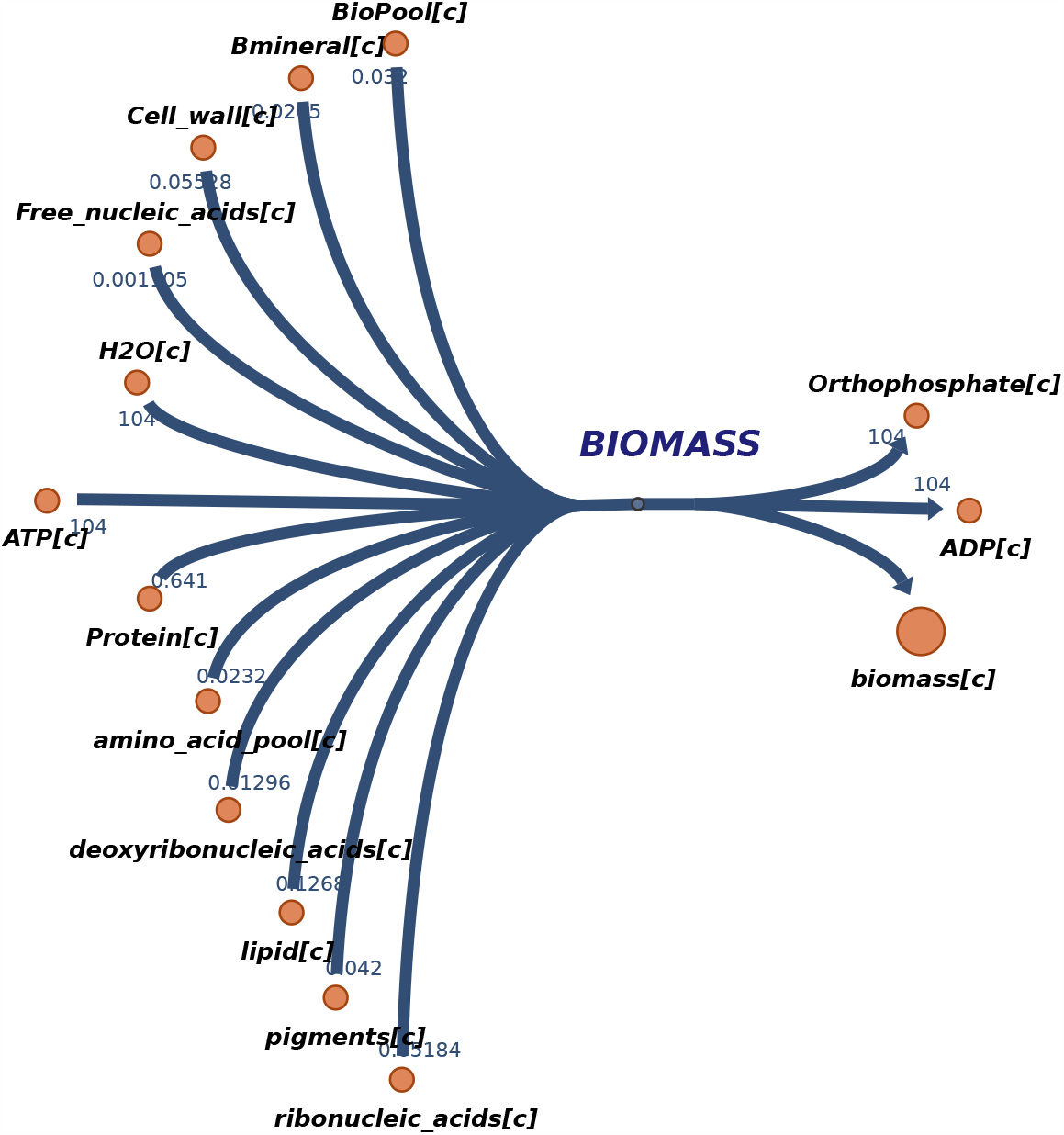
Biomass reaction of *Prochloroccocus MED4* in the model iSO595 (produced with Escher)

To compute an overproduction of those components, we created exchanges reactions allowing the organism to produce them without necessarily using them for biomass production. Those exchange reactions were restricted to production only (lower bound of 0). Hence, these changes in the metabolic network do not change the growth computed through our formalism but enables us to compute the overproduction of these metabolites, allowing us to investigate the organism’s physiology and step away from the fixed stoichiometry imposed by the metabolic framework. Further investigation is needed to fully allow the GSM to control its biomass composition.

### 1.C. Resource constraint interpretations

The resource constraint can be linked to a metabolic stress measure. For instance, in a strongly constrained environment (resource constraint near 100%), the growth is assumably limited by the considered nutrient.

Moreover, the resource constraint can be used with a growth allocation metric to further study the modeled organism. The growth allocation for nutrient *i* would be defined as the ratio of the actual uptake flux of nutrient over the bioavailability of the nutrient 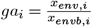, where 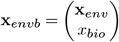 and 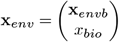 as defined in Material & Methods. Hence, a growth allocation of 1 is equivalent to a constraint of 100%, meaning that all the available nutrients are used to produce growth. Lower values of growth allocation can be compared to the associated resource constraint to see the percentage of used nutrients for growth (growth allocation) and the available nutrient for auxiliary metabolism (resource constraint).

### 1.D. DOC metabolites

**Table 1.** Lists of metabolites that contribute to Dissolved Organic Carbon for each GSM, with their stoichiometric coefficient for carbon.

| *Prochlorococcus MED4* | *Thalassiosira pseudonana* |
| --- | --- |
| L-Arginine (6), L-Asparagine (4), L-Aspartate (4), L-Glutamate (5), L-Glutamine (5), Glycine (2), L-Histidine (6), L-Isoleucine (6), L-Leucine (6), L-Lysine (6), L-Methionine (5), L-Phenylalanine (9), L-Proline (5), L-Serine (3), L-Threonine (4), L-Tryptophan (1), L-Tyrosine (9), L-Valine (5), D-Glucose (6), Acetate (2), Citrate (6), Glutathione (10), Pantothenate (9), Succinate (4), Thymidine (10), Xanthosine (10), 4-Aminobenzoate (7), 5-Methylthioadenosine (11), Adenine (5), Guanine (5), Guanosine (10), Putrescine (4), Spermidine (7), 4-Hydroxybenzoate (7), S-Malate (4) | Folate (19), N-acetyltaurine (4), Acetate (2), Choline (5), Uracil (4), Xanthine (5), L-Glutamate (5), L-Aspartate (4), L-Isoleucine (6), L-Leucine (6), L-Valine (5), L-Asparagine (4), L-Alanine (3), L-Glutamine (5), L-Histidine (6), L-Serine (3), L-Threonine (4), Glycine (2), L-Proline (5), Adenosine monophosphate (10), 3-(dimethylsulfonio)propanoate (5) |

## Footnotes

1 Thiele, I. & Palsson, B. Ø. A protocol for generating a high-quality genome-scale metabolic reconstruction. Nat Protoc 5, 93–121 (2010).

2 Varma, A. & Palsson, B. O. Metabolic Flux Balancing: Basic Concepts, Scientific and Practical Use. Nat Biotechnol 12, 994–998 (1994).

3 Srinivasan, S., Cluett, W. R. & Mahadevan, R. Constructing kinetic models of metabolism at genome-scales: A review. Biotechnology Journal 10, 1345–1359 (2015).

4 Ofaim, S., Sulheim, S., Almaas, E., Sher, D. & Segrè, D. Dynamic Allocation of Carbon Storage and Nutrient-Dependent Exudation in a Revised GenomeScale Model of Prochlorococcus. Front. Genet. 12, (2021).

